# Molecular Response to PARP1 Inhibition in Ovarian Cancer Cells as Determined by Mass Spectrometry Based Proteomics

**DOI:** 10.1101/2021.03.12.435005

**Authors:** Alexandra Franz, Fabian Coscia, Ciyue Shen, Lea Charaoui, Matthias Mann, Chris Sander

**Author notes:** Correspondence to (AF), (CS).

## Abstract

Poly (ADP)-ribose polymerase (PARP) inhibitors have entered routine clinical practice for the treatment of high-grade serous ovarian cancer (HGSOC), yet the molecular mechanisms underlying treatment response to PARP1 inhibition (PARP1i) are not fully understood. Here, we used unbiased mass spectrometry based proteomics with data-driven protein network analysis to systematically characterize how HGSOC cells respond to PARP1i treatment. We found that PARP1i leads to pronounced proteomic changes in a diverse set of cellular processes in HGSOC cancer cells, consistent with transcript changes in an independent perturbation dataset. We interpret decreases in the levels of the pro-proliferative transcription factors SP1 and β-catenin and in growth factor signaling as reflecting the anti-proliferative effect of PARP1i; and the strong activation of pro-survival processes NF-ϰB signaling and lipid metabolism as PARPi-induced adaptive resistance mechanisms. Based on these observations, we nominate several protein targets for therapeutic inhibition in combination with PARP1i. When tested experimentally, the combination of PARPi with an inhibitor of fatty acid synthase (TVB-2640) has a 3-fold synergistic effect and is therefore of particular pre-clinical interest. Our study improves the current understanding of PARP1 function, highlights the potential that the anti-tumor efficacy of PARP1i may not only rely on DNA damage repair mechanisms and informs on the rational design of PARP1i combination therapies in ovarian cancer.

## Background

High-grade serous ovarian cancer (HGSOC) is the most common and one of the most aggressive subtype of ovarian cancer, responsible for the majority of ovarian cancer deaths (1). A crucial step forward for the treatment of HGSOC patients was the introduction of molecular inhibitors that target poly (adenosine diphosphate (ADP)-ribose) polymerase (PARP) enzymes (2). PARP enzymes play central roles in the DNA damage response and their inhibition has demonstrated significant improved progression-free survival in HGSOC patients that lack homologous recombination (HR) capacity through mutations in *BRCA1* or *BRCA2* genes (3,4). Three PARP inhibitors (PARPi) have received regulatory approval by the U.S. Food and Drug Administration (FDA) in specific clinical settings (3,5–8). Such as, PARPi have been approved as maintenance therapy for ovarian cancer patients with a germline or somatic BRCA1/2 mutation after completion of adjuvant treatment. PARP inhibitors are also used in the recurrent setting, especially in patients that had not been treated with PARP inhibitors. Other indications for treatment with PARPi include patients with mutations in other HR pathway components (e.g., *BRIP1*, *PALB2*, *FANCD2*) (9).

Several models have been put forward to explain the mechanisms of PARPi action, but understanding is still incomplete (10). Originally, it was widely believed that the therapeutic benefit of PARPi is primarily due to unrepaired single-stranded DNA breaks, which are converted to irreparable toxic DNA double-strand breaks (DSB) during replication in HR-deficient cells and lead to cell death (11). Other models suggest that due to the diverse roles of PARP proteins in the DNA repair response, PARPi-induced deficiencies in other DNA repair pathways may also contribute to tumor sensitivity to PARPi (12). A more recent model is the ‘trapping’ of PARP proteins at DNA damage sites during DNA replication, leading to replication collapse, the accumulation of unresolved DSBs and eventually to cell death (13,14).

Beyond its primary roles in DNA repair, the PARP1 protein, which is responsible for over 80% of PARP enzyme-related activities (15), has also been shown to directly bind and co-activate oncogenic transcription factors that promote cell proliferation, growth and differentiation in different human cancer models (16). For example, in the context of colorectal cancer, it was demonstrated that PARP1 binds transcription factor TCF4 and interacts in a complex with TCF4 and β-catenin contributing to early colorectal carcinogenesis (17). In osteosarcoma tumor cells, PARP1 has been found to co-activate oncogenic transcription factor B-MYB (18). Similarly, in a model of human skin cancer and chronic myelogenous leukemia, it was shown that PARP1 positively regulates the activity of activator protein-1 and hypoxia inducible factor 1 alpha, thereby promoting tumor cell survival and tumor growth (19). Also, several studies showed that PARP1 directly interacts with transcription factor nuclear factor kappa B (NF-ϰB) and functions as its transcriptional co-activator for ϰB-dependent gene expression (20). However, although additional proteins have been reported to interact with PARP1, little is known about their relevance for the anti-tumor activity or the emergence of resistance to PARP1i in the context of HGSOC (10,11). Indeed, despite their clinical efficacy in HGSOC with deficiencies in HR, PARP inhibitors have also been found to be effective in HGSOC patients without *BRCA* loss-of function mutations or homologous recombination deficiency as determined by last generation genomic sequencing (3,4) and the FDA and EMA (European Medicines Agency) have recently broadened the approval of PARPi as maintenance monotherapy for ovarian cancer patients independently of *BRCA* or HR status (21). Of note, alterations in *RAD51C*, *RAD51D* and high-level *BRCA1* promoter methylations were recently associated with positive response to PARPi rucaparib suggesting broadening genomic testing (22). An improved understanding of the cellular mechanisms underlying PARPi treatment response is thus needed to allow more accurate treatment decisions involving PARPi therapies as well as to overcome often seen PARPi resistance (23).

Here, we set out to comprehensively assess the cellular response PARP1i in HGSOC cells using quantitative mass spectrometry (MS)-based proteomics and data-driven protein network analysis. To this end, we systematically analyzed the proteomic response profile following PARP1i treatment to 1) explore PARP1’s role in cellular processes beyond the DNA damage response in HGSOC cells, 2) identify potential markers of PARP1i sensitivity or resistance and 3) determine candidate targets for PARP1i combination therapies.

## Results

### PARP1i perturbation profiling in Ovsaho ovarian cancer cells using MS-based proteomics

Protein response profiling allows one to monitor and capture cellular behavior in response to drug perturbation providing information about molecular processes involved in treatment response that contribute to sensitivity or resistance (24,25). Thus, to gain insight into the molecular processes induced by the inhibition of PARP1 protein in HGSOC, we treated HGSOC Ovsaho cells with a potent inhibitor AG-14361 that selectively inhibits the PARP1 protein (Ki < 5nM) (26), and measured the protein response profile by unbiased MS based-proteomics. We selected Ovsaho cells, which are a well-defined preclinical model of HGSOC reflecting the genomic and proteomic features of HGSOC patient tumors (27,28). Ovsaho cells carry a copy number deletion of the *BRCA2* gene and mutations in the *FANCD2* gene, which is another component of the HR pathway (Figure 1A) (29). We treated Ovsaho cells at the drug’s half-maximal inhibitory concentration (IC50) to inhibit cell proliferation by 50% after 72 hrs as determined by dose-response curves. The treatment time was set to 72hrs to measure what is plausibly the steady state adaptive effect of the cells on PARP1 inhibition (PARP1i). Ovsaho cells were treated with increasing concentrations of PARP1 inhibitor AG-14361 and cell viability was examined for 72 hrs by counting cells using live-imaging (Material and Methods). Treatment with PARP1 inhibitor resulted in a concentration-dependent decrease of cell numbers with IC50 value at 20 *μ*M after 72 hrs (Figure 1B).

**Figure 1.**
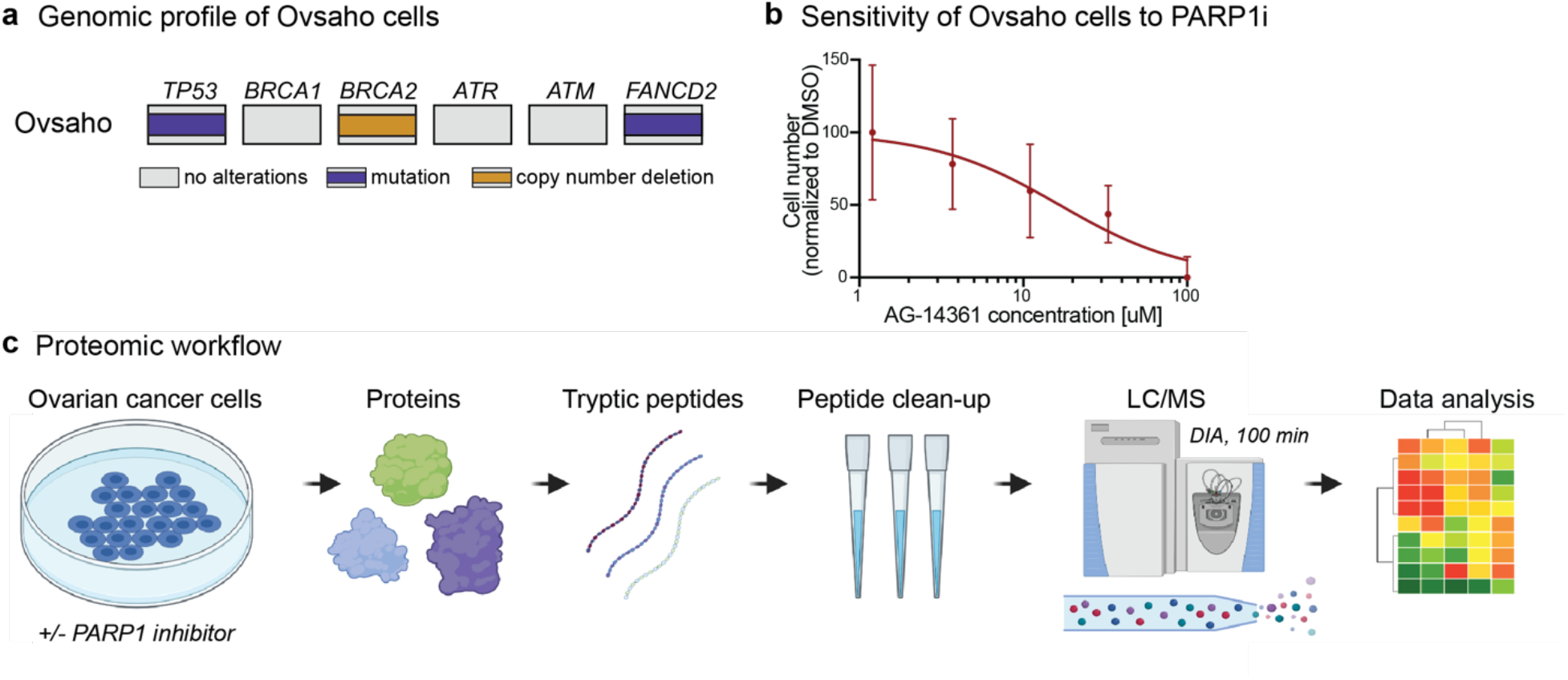
Measuring the sensitivity to PARP1 inhibition in Ovsaho cells and drug response profiling using LC/MS-MS-based proteomics. (a) Genomic profile of Ovsaho cells with HR-associated genes. Ovsaho cells have a mutation in the *TP53* and *FANCD2* gene and a copy number deletion of *BRCA2* gene. (b) Dose-response curve based on cell number count after 72hr. PARP1 inhibitor AG-14361 treated Ovsaho cells versus control treatment. DMSO was used as control vehicle. Data represent three independent experiments and error bars represent standard derivation of technical replicates with a total number of experiments = 9. (c) Schematic view of the LC/MS-MS workflow. PARP1i-treated Ovsaho ovarian cancer cells were prepared for LC/MS-MS. Following protein extraction and tryptic digest, proteins were separated and measured in single runs using a quadrupole Orbitrap mass spectrometer. Label-free protein quantification was performed using the Spectronaut software environment. (LC-MS) Liquid Chromatography with tandem mass spectrometry. Figure 1C was created with BioRender.com.

Next, Ovsaho cells were treated at inhibitor IC50 concentration (20*μ*M) or DMSO as control in three biological replicates and samples were collected after 72 hrs for MS-based proteomics (Figure 1C). We employed a data-independent acquisition (DIA) method combined with label-free based quantification (Material and Methods), resulting in ~5,000 quantified protein groups per single replicate measurement (Figure 2A). Biological replicates of both PARP1i-treated and control cells correlated well with median Pearson correlation coefficients of 0.91 and 0.95, respectively (SFigure 1A). We evaluated replicate reproducibility by calculating the coefficient of variation (CV) between biological replicates. The results of the analysis revealed a median CV of less than 20% between biological replicate protein measurements confirming good experimental reproducibility between replicates (SFigure 1B).

**Figure 2.**
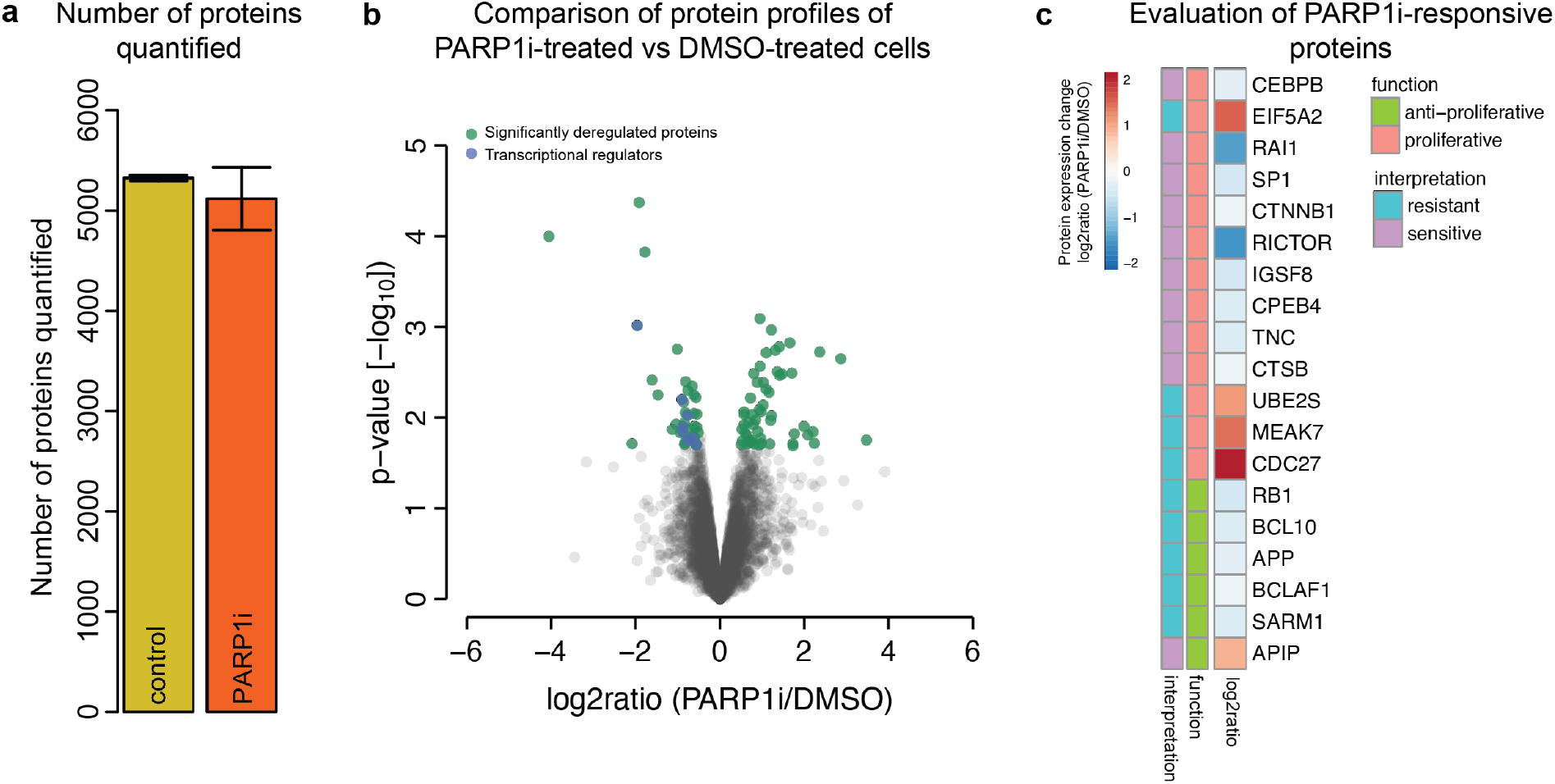
PARP1 inhibitor induced protein response profile in Ovsaho cells. (a) Number of protein groups detected by MS for each perturbation condition, three biological replicates per condition. Numbers are reported as mean of the three biological replicates and error bars show standard deviation for each condition. (b) Identification of PARP1i induced proteins whose expression is significantly changed compared to control DMSO treatment. Volcano plot of statistical significance against log2 protein expression change between PARP1i-treated versus control cells after 72hrs. In green are significantly expressed proteins with log2ratio (PARP1i/DMSO) ≤ −0.5 or ≥ 0.5 with p-value ≤ 0.05 (FDR < 0.2) and in blue proteins with transcriptional activity (STable1). (c) PARP1i-responsive proteins involved in pro-roliferative or anti-proliferative processes and possible interpretation for their relevance to sensitivity and resistance to PARP1i based on their proliferative function and protein expression change level upon PARP1i treatment relative to control (DMSO) treatment. If a pro-proliferative protein (or a protein with pro-proliferative functions) was upregulated upon PARPi treatment, it was considered as marker of resistance to PARPi; if it was downregulated, it was considered sensitive to PARPi. The reverse holds for the anti-proliferative proteins.

### Identification of PARP1i-responsive proteins

Based on our reproducible mass spectrometric measurements, we next analyzed the proteomic alterations after PARP1i treatment. We first identified which proteins were significantly changed upon PARP1i treatment by comparing the protein expression levels values of PARP1i-treated samples to control samples. We filtered for proteins with a quantitative difference of at least 1.41-fold (log2ratio (PARP1i/DMSO) > 0.5 or < −0.5). Statistical significance of the protein expression level change was determined by statistical testing using an unpaired t-test with a p-value of < 0.05 providing statistical evidence that there is less than 5% probability that the change in expression is due to natural random variation. In addition, we applied Benjamini-Hochberg (BH) correction for multiple hypothesis testing to control the false discovery rate (FDR) (30,31) (Material and Methods).

Overall, we identified 93 significantly regulated proteins after PARP1i treatment relative to control treatment of which 54 were increased, and 39 proteins decreased (Figure 2B, STable1).

Of note, these included several proteins with well-known associations to PARP1 in ovarian cancer. For example, inhibition of PARP1 affected the expression of proteins involved in DNA repair mechanisms including DNA repair scaffold protein XCCR1 (X-ray cross complementing protein 1) (log2ratio (PARP1i/DMSO) = −1.037, p-value = 0.01) and DNA damage regulator TTI2 (Tel two-interacting protein 2) (log2ratio (PARP1i/DMSO) = 0.69, p-value = 0.01). We also identified proteins that are not part of the DNA repair response and to our knowledge have not yet been reported in the context of PARPi-treated HGSOC cells (STable1). Most notably, we found that treatment with PARP1i significantly altered the expression level of proteins with transcription- and translation-regulatory activities, including transcription factors SP1 (specificity protein 1), RB1 (retinoblastoma protein 1), CPEB4 (cytoplasmic polyadenylation element binding protein 4), CEBPB (CCAAT enhancer binding protein beta), RAI1 (retinoic acid-induced gene 1), β-catenin and BCL10 (B cell lymphoma 10) (Figure 2B, C, STable1). Since these proteins are well known to be involved in diverse cellular processes by regulating the transcription of a variety of genes (32–38), we next assessed their potential relevance to PARP1i-induced anti-tumor effects or resistance mechanisms.

### Individual PARP1i-responsive proteins as markers of PARP1i sensitivity and resistance

To analyze whether our identified PARP1i-responsive proteins are associated with treatment sensitivity or resistance, we inferred their anti- and pro-proliferative properties based on a review of the literature research and assessed their protein expression level change upon PARP1i. Our analysis revealed several proteins that have been reported to promote cell proliferation (STable1). Moreover, we found that the majority of these proteins were significantly downregulated upon PARP1i treatment. For example, we found a marked decrease of transcription factors β-catenin (log2ratio (PARP1i/DMSO) = −0.56; p-value = 0.01) and SP1 (log2ratio (PARP1i/DMSO) = −0.88; p-value = 0.01) (Figure 2C), which induce the expression of genes involved in the promotion of cell proliferation such as *c-myc* and *CCND1* (32,36). The broad PARP1i-induced downregulation of pro-proliferative transcription factors could thus possibly contribute to PARP1i-induced cytotoxicity.

We also identified proteins that could contribute to adaptive survival mechanisms in response to PARP1i (Figure 2C). For example, we found that tumor suppressor RB1 was strongly downregulated (log2ratio (PARP1i/DMSO) = −0.88; p-value = 0.01), whereas candidate oncogene eIF5A2 (eukaryotic translation initiation factor 5A isoform 2) was strongly upregulated (log2ratio (PARP1i/DMSO) = 1.411; p-value = 0.001). Similarly, cell cycle promoting factors ubiquitin-conjugating enzyme E2S (UBE2S, (log2ratio (PARP1i/DMSO) = 0.932; p-value = 0.0081) and CDC27 (cell division cycle 27; log2ratio (PARP1i/DMSO) = 1.99; p-value = 0.012) showed a marked increase of protein expression levels upon PARP1i. In sum, these observations show that HGSOC Ovsaho cancer cells reprogram a diverse array of proteins in response to PARP1i treatment, of which some could plausibly contribute to PARP1i-induced cytotoxicity or resistance due to their reported anti- and pro-proliferative properties.

### Identification of functional protein networks induced by PARP1i

Understanding the impact of a drug on the protein network can provide further insight into the context in which the drug target works. Therefore, we next aimed to identify the cellular context of PARP1i-responsive proteins. To this end, we performed protein network analysis using the data-driven computational framework NetboxR, which identifies functional protein network modules independent and across the boundaries of pre-defined protein lists by integrating protein network interactions with a clustering algorithm. This data-driven approach thus allows the discovery of *de novo* functional protein network modules and overcomes the issue of the occurrence of proteins in more than one of the curated protein/gene sets (39,40) (Material and Methods). We annotated identified protein network modules with pathway process labels by assigning module protein members to curated pathways based on the Reactome database using the pathway enrichment analysis tool g:profiler with an adjusted (adj.) p-value < 0.05 threshold (41).

Our data-driven protein network analysis revealed 23 functional protein network modules comprising 117 proteins that significantly respond to PARP1i (FDR-corrected p-value < 0.05) (STable2). Figure 3A shows the effects of PARP1i treatment on major protein networks. In this figure, each node represents a protein and edges represent protein interactions. Our analysis revealed several processes that have previously been reported to be associated with PARPi treatment in ovarian cancer cells, confirming our approach’s ability to identify molecular processes underlying PARP1i treatment responses. For example, we found protein modules whose proteins were significantly over-represented in cellular senescence (adj. p-value = 1.025×10^−4^), DNA repair (adj. p-value = 1.882×10^−2^) and G2-G2/M cell cycle phase (adj. p-value = 7.014×10^−4^) (Figure 3A, B; STable2) (42,43). In addition, we identified one protein module annotated as MAPK activation (adj. p-value = 2.247×10^−3^) (STable2). MAPK activation has previously been identified as PARPi resistance mechanism and has been reported as a strong candidate for PARP1i combination therapy (44). Consistent with our results on the individual protein level, our protein network analysis also revealed PARP1i-responsive processes with reported pro-proliferative properties that have not yet been reported in association with PARP1 in HGSOC cells, such as NF-ϰB signaling (adj. p-value = 3.456×10^−2^), EGFR signaling (adj. p-value = 3.421×10^−3^) and lipid metabolism (adj. p-value = 1.197×10^−6^) (Figure 3A, B; STable2). We therefore next asked whether these processes could likely contribute to PARP1i-induced cytotoxicity or resistance.

**Figure 3.**
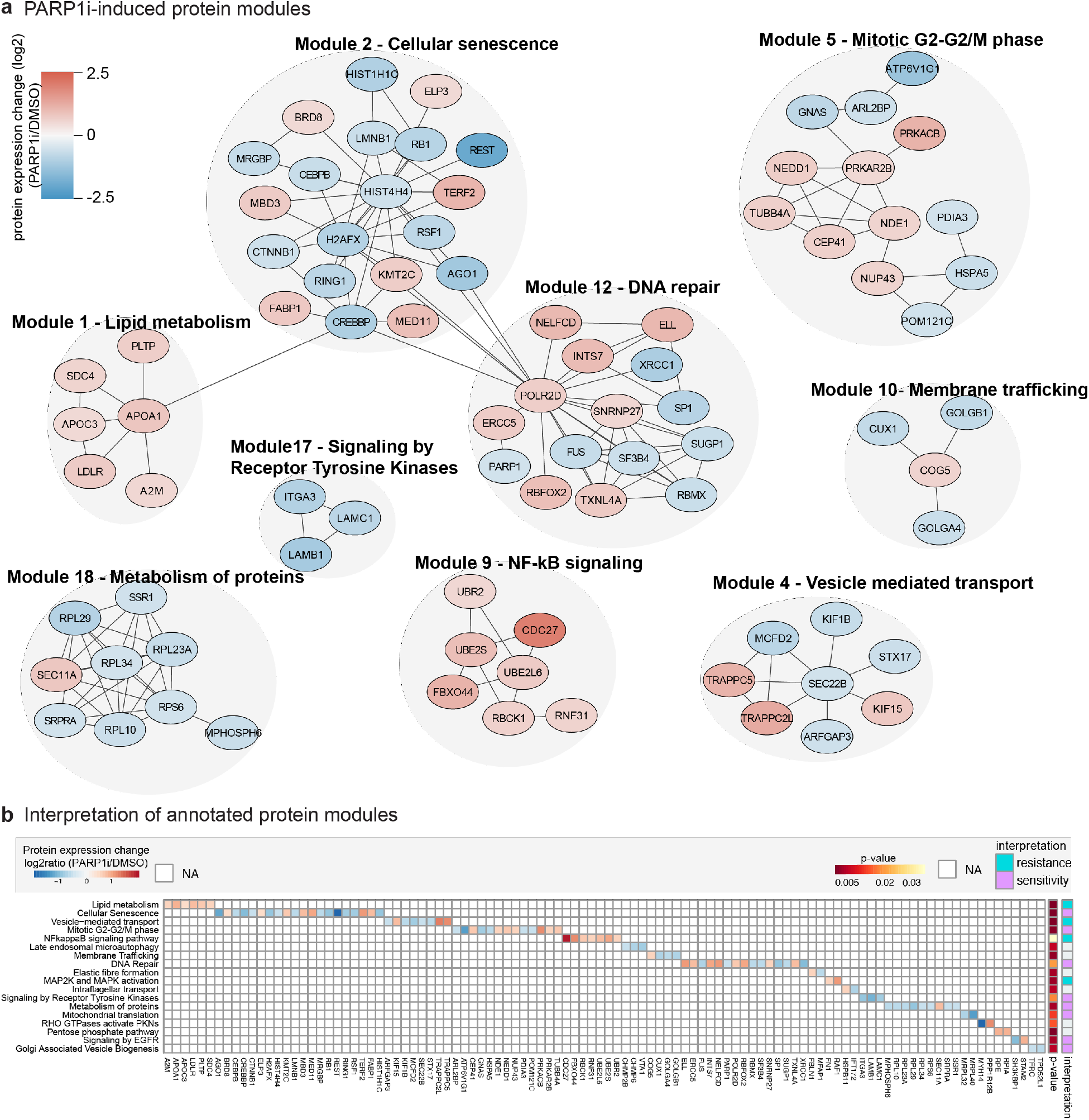
Data-driven exploration of protein signaling network modules in response to PARP1i. Proteins were grouped into functional modules using Netbox algorithm (FDR corrected p-value ≤ 0.05), which combines prior-knowledge of protein network interactions with a clustering algorithm to identify functional protein modules across the boundaries of curated and pre-defined lists of proteins (28,29) (Material and Methods). As a next step, protein modules were characterized by assigning module protein members to pathways based on the Reactome database using g:profiler analysis (adj. p-value ≤ 0.05) (STable2). (a) Depicted are annotated protein modules with > 3 protein members. Nodes represent proteins and are colored based on protein expression change (log2ratio PARP1i/DMSO). Edges represent protein interactions. (b) Heatmap showing annotated protein modules and protein expression changes of corresponding protein members (STable2). Based on their function and protein expression changes, annotated protein modules were evaluated to be associated with PARP1i-induced sensitivity or resistance (STable2).

### Molecular processes associated with PARP1i cytotoxicity or resistance

Based on the function of identified protein network modules in proliferative processes and the protein expression change of module members, we evaluated whether PARP1i-responsive processes could promote cytotoxicity or resistance. Most notable, we found that the inhibition of PARP1 leads to an overall decreased activity of cell growth-inducing signaling pathways, including tyrosine kinase receptor signaling (adj. p-value = 1.680×10^−2^) and EGFR signaling (adj. p-value = 3.421×10^−3^), suggesting that these processes could contribute to PARP1i-induced cytotoxicity (Figure 3B). We also discovered cellular networks that are indicative of adaptive response mechanisms to PARP1i. In particular, we noted a large cluster of nodes related to NF-ϰB signaling (adj. p-value 3.456×10^−2^) and lipid metabolism (adj. p-value 1.197×10^−6^) with all protein members being highly expressed in response to PARP1i (Figure 3A, B). NF-ϰB is a pivotal factor in regulating immune responses and inflammation and has been implicated in tumor growth. In ovarian cancer, it was found that the activation of NF-ϰB promotes tumor cell proliferation (44). Moreover, activated NF-ϰB signaling has been observed in response to diverse anti-cancer therapeutic agents in several pre-clinical studies and its inhibition has been reported to enhance apoptosis and to re-sensitize drug resistant tumors potentiating the anti-tumor efficacy of anti-cancer therapies (45,46). Furthermore, a more recent study reported high levels of NF-ϰB activity in PARPi-resistant human ovarian cancer cell line UWB1.289, which derived from a tumor of papillary serous histology. The authors demonstrated that inhibition of NF-ϰB with inhibitor bortezomib increased the sensitivity to PARPi olaparib in PARPi-resistant UWB1.289 cells (47). Similarly, increased lipid metabolism is involved in chemotherapy resistance and other small agents such as VEGF neutralizing antibody bevacizumab. The inhibition of lipid metabolism enzymes has been shown to decrease tumor growth (48). Based on these observations, we interpret the increased protein expression levels of proteins involved in NF-ϰB signaling and lipid metabolism in response to PARP1i treatment as adaptive mechanism of cells to survive, while reduced growth factor signaling and reduced mitochondrial translation might contribute to cytotoxicity.

### Confirmation of PARP1i-induced cellular processes in other HGSOC cell lines

We next validated our findings using independent perturbation datasets involving different HGSOC cell lines and PARPi olaparib and rucaparib. To this end, we employed the recently published L1000 dataset from the LINCS (library of integrated network-based cellular signatures) data portal. The L1000 is a gene expression assay in which the mRNA expression profiles of specific 978 landmark genes are measured in response to drug perturbations (49). From the dataset, we found three cell lines TYKNU, COV644 and RMUGS that closely match patient HGSOC tumors (27) and were treated with pan PARP1/2 inhibitors olaparib and rucaparib. By applying data-driven protein network analysis on strongly drug responsive genes (log2ratio (inhibitor/control) > 0.5 or < −0.5) (Material and Methods) and annotating identified functional modules with pathway labels using pathway enrichment (Material and Methods), we re-captured almost all previously identified annotated protein networks across tested cell lines (FDR corrected p-value < 0.05), including tyrosine kinase signaling, mitochondrial translation, NF-κB signaling and lipid metabolism (Figure 4 A,B: STable3). Correlation analysis of the overall protein module activity based on overlapping annotated protein network modules between L1000 and our proteome profiles (Material and Methods), revealed a positive correlation of olaparib-induced module activity with PARP1i-induced module activity (Pearson correlation coefficient R = 0.37; Material and Methods, STable3, SFigure2) with the olaparib-treated cell line RMUGS showing the highest correlation in activity (R = 0.83, Figure 4C).

**Figure 4.**
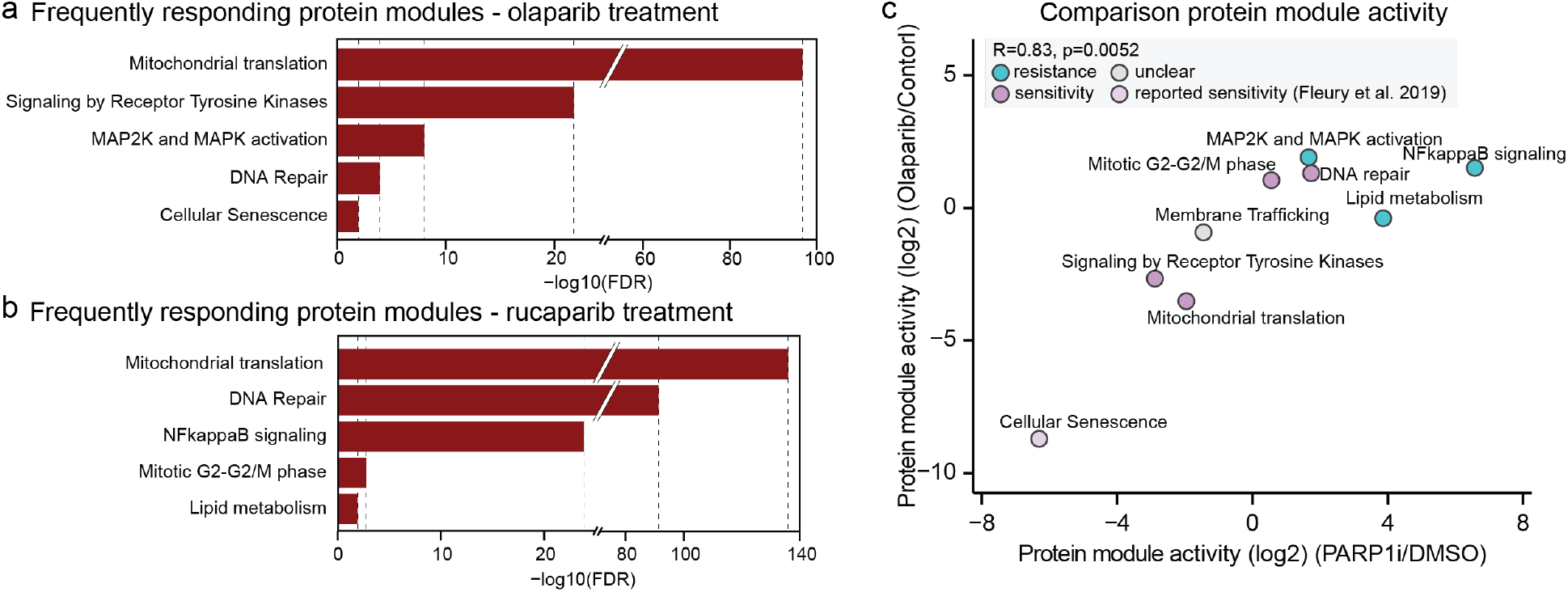
Confirmation of PARP1i-induced protein networks in HGSOC cell lines. Protein network and pathway analysis were performed on independent L1000 transcriptome dataset for HGSOC cell lines RMUGS, COV644 and TYKNU treated with pan-PARPi olaparib and rucaparib as described previously (FDR-corrected p-value < 0.05; Material and Methods, STable3). (a, b) Barplots represents top 5 frequently responding annotated protein modules (adj. p-value < 0.05) across all cell lines (STable3). (c) Correlation analysis of the overall protein module activity based on overlapping annotated modules between L1000 olaparib-treated RMUGS cell line and proteome profiling. Activity was calculated by integrating protein expression changes in modules (38) (Material and Methods).

However, comparing PARP1i-induced module activity to those induced by rucaparib treatments revealed a negative correlation (R = −0.52; Material and Methods, STable3, SFigure2). This might be related to the different specificities of PAPRPi and rucaparib for the PARP1 protein (50).

### Synergistic PARP1i drug combinations in Ovsaho ovarian cancer cells

Strong protein signals in response to drug perturbations provide a strong basis for the rational design of combination therapies (24,25). Based on our observations on protein expression and functional protein networks, we nominate four promising targets as candidates for PARPi combination studies. These candidate combination targets had the strongest activation upon PARP1i treatment and have been previously associated to promote pro-proliferative processes (47,48,51,52) (Table 1).

**Table1.** Nomination of candidate PARPi combination therapies based on identified molecular processes and existing small molecule drugs with clinical relevance

| Molecular process | Target | Drug combination with PARPi |
| --- | --- | --- |
| Lipid metabolism | FASN | PARPi + TVB-2640 |
| NF-κB signaling | IκBα | PARPi + Bortezomib |
| Cell proliferation | eIF5A-2 | PARPi + N(1)-guanyl-1,7,-diamineoheptane (GC7) |
| Mitotic exit | UBE2S/APC/C | PARPi + proTAME |

We therefore experimentally tested inhibition of these targets in combination treatment with PARP1i AG-14361 in Ovsaho ovarian cancer cells and examined if the combination treatment could enhance the efficacy of PARP1i. To this end, we assessed cell viability of Ovsaho cells to PARP1i combination and single agent treatments in dose-response curves for 72 hrs by live-imaging using the IncuCyte Zoom system (Material and Methods). To quantify drug synergy (Combination Index, CI) we used the Chou-Talaly method (53). In case several inhibitors were available for a specific target, we selected a compound with reported clinical relevance (48,51,54,55) (Table 1). The combination treatments revealed synergistic, other non-additive or additive effects after 72hrs at IC50 as follows (Figure 5). PARP1i combination with TVB-2640, an inhibitor of fatty acid synthase (FASN) (56) has a strong synergistic effect (CI = 0.35), while PARP1i combination with GC7, an inhibitor of eIF5A-2, has a smaller synergistic effect (CI = 0.64) (Figure 5). Bortezomib, which inhibits the proteasome and the NF-kB signaling pathway (47), strongly decreases cell viability as a single agent and at the same concentration is more effective than single PARP1i treatment. Combination treatment by both inhibitors at the same total concentration as that of the single drugs has a less than additive effect (CI = 1.7). The combination treatment by PARPi with APC inhibitor proTAME (54) is additive (with CI = 0.95, Figure 5).

**Figure 5.**
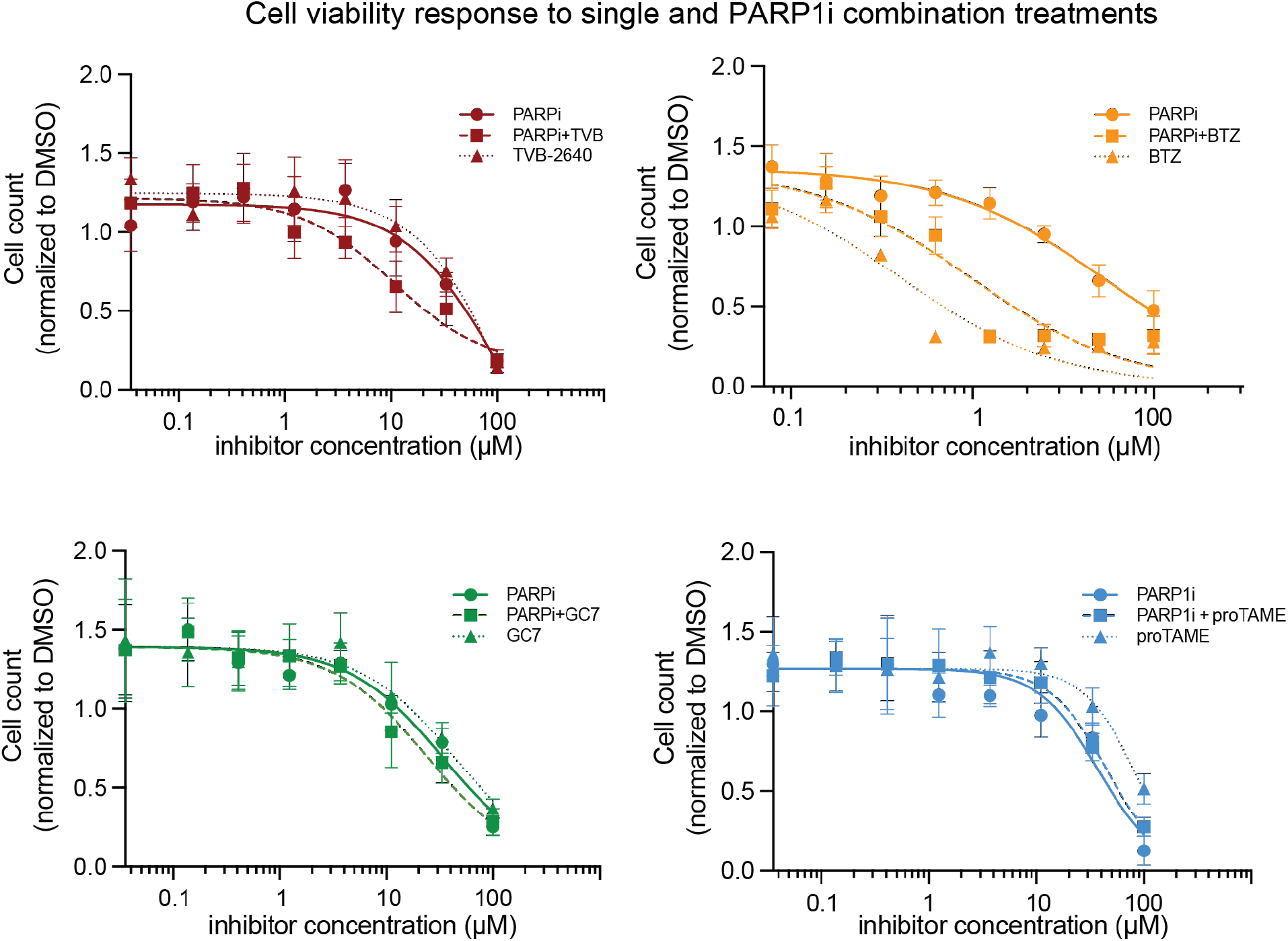
Effect of PARP1i drug combinations in Ovsaho ovarian cancer cells. Dose-response curve of cell viability based on cell number count for PARP1i AG-14361 + TVB-2640 (top left), PARP1i AG-14361 + Bortezomib (top right), PARP1i AG-14361 + proTAME (bottom left) and PARP1i AG-14361+ GC7 (top right) after 72hr. Inhibitor concentrations were combined in 1:1 ratio. DMSO was used as control vehicle. Data is representative of three independent experiments for treatment and error bars are the standard deviation of technical replicates for n= 9. Cell counts were measured using live-im- aging in an IncuCyte Zoom microscopic station (Material and Methods). (BTZ) Bortezomib, (TVB) TVB-2640.

Based on these results, we nominate three of the four identified PARP1i drug combinations for future pre-clinical investigations.

## Discussion

It is widely believed that the anti-tumor efficacy of PARPi is due to synthetic lethality between PARPi and defects in HR deficient cancer cells (11). However, PARPi’s have also demonstrated therapeutic benefits in HGSOC patients regardless of HR status (21). In addition, several studies report that the PARP1 protein is implicated in cell growth and cell proliferation by modulating the activity of transcription factors (16), which leads to the notion that PARPi might also confer antitumor effects beyond the DNA repair mechanism and indicate a broader utility for PARPi in the treatment of cancer patients (10).

Our proteomics data support this view. In agreement with previous studies (16), we find that PARP1i blocks protein expression of many pro-proliferative proteins; in particular, transcriptional activators, which plausibly contribute to PARP1i-induced cytotoxicity in our HGSOC model system. However, the roles of PARP1 as transcriptional co-activator or repressor of transcription factors are known to be complex, gene and context-specific (57). Further studies are thus warranted to precisely delineate the relationship of PARP1 with individual pro-proliferative transcription factors and the effect on the expression of target genes and contribution to cell proliferation in HGSOC cells.

PARPi offers a significant clinical benefit, but resistance to PARPi typically develops in most advanced ovarian cancers (58). Most of the mechanisms of PARPi resistance identified involve the restoration of stalled replication fork protection, as well as reactivation of the HR pathway based on genetic and epigenetic changes in the tumor (59). Frequently reported is for example the restoration of functional BRCA1/2 through secondary mutations, BRCA1 promoter methylation or the rewiring of the DNA damage response network (23). Therefore, multiple strategies to overcome PARPi resistance have been designed to selectively disrupt HR in cancer cells, e.g., via combined inhibition of PARP and CDK1 (60), and thus to re-sensitize the cells to PARPi (59). Our unbiased proteomics approach reveals other potential resistance mechanisms to PARPi treatment. The combination of PARPi with the inhibitor of FASN (TVB-2640) has an approximately 3-fold synergistic effect. There is evidence that lipid metabolic reprogramming can mediate resistance to anti-cancer agents including chemotherapy and targeted therapies including in breast, lung and prostate cancer cell lines and xenograft models (61). Indeed, targeting altered metabolic lipid pathways with pharmacologic interventions has become an emerging anti-tumor strategy in several cancer types with currently 10 inhibitors being tested in clinical trials including TVB-2640 (62). While in ovarian cancer overexpressed lipogenic enzymes are increasingly seen as potential therapeutic opportunity to inhibit cancer growth, peritoneal metastasis and/or overcome resistance to chemotherapy or anti-angiogenic therapy (61,63,64), our study is the first demonstration that the combined inhibition of PARP1 and FSAN represents an synergistic anti-proliferative combination strategy. FASN has been reported to function as a central regulator of the lipid metabolism, but it has also been suggested that FASN is involved in glycolysis and amino acid metabolism (65). Therefore, we cannot fully exclude the possibility of the contribution of pathways other than lipid metabolism, such as glycolysis, to the observed synergistic effect. Additional studies with small-molecule inhibitors that more specifically target metabolic lipid pathways are thus needed to gain further mechanistic insight.

Our drug combination assay also shows a synergistic effect of the dual inhibition of PARP1 and UBE2S (see Figure 5). UBE2S has been reported to induce mitotic exit (52). Since mitotic exit by slippage has been found as typical response of *TP53*-mutant cancer cells to resist drug-induced mitotic cell death and blocking mitotic exit has already shown to successfully reduce tumor growth in paclitaxel-treated ovarian cancer models (54), we speculate that additional blocking of mitotic exit by APC/C inhibitor proTAME could have enhanced the anti-tumor effects of PARP1i. However, additional experiments will be important to clarify the mechanisms underlying the enhanced cytotoxic effects of inhibition of UBE2S.

We find that BTZ single agent treatment is strongly cytotoxic in Ovsaho cells and is more effective than single PARP1i treatment. Therefore, it is not surprising that the combination treatment by BTZ and PARP1i using the same total concentration has a weaker effect than BTZ alone. Optimization of concentration ratio at which these two drugs could be usefully combined will be important in follow-up studies. Indeed, a recent study found that PARPi and BTZ combination treatment at a different concentration ratio than the one used here was more effective than single agent treatments in the chemosensitive ovarian cancer cell line OV2008 (66); and another study demonstrated the effectiveness of BTZ treatment in PARPi-resistant cells (47).

Our drug combination results are promising with evidence of additive or synergistic effects. However, our study is limited to the Ovsaho HGSOC cell line and AG-14361 PARP1i inhibitor. Additional studies in a diverse panel of HGSOC cell lines, organoids and mouse models, using FDA-approved panPARP1/2 inhibitors (olaparib, rucaparib and niraparib) will thus be important to explore the therapeutic potential of the combinations and determine differential sensitivity. Even non-synergistic combinations might be useful clinically, either by reducing the likelihood of the emergence of resistance or by a higher probability of therapeutic benefit in a patient population with genetically heterogeneous and differentially sensitive tumors (67,68). Another important aspect for clinical translation is to examine these drug effects in HGSOC model systems with different BRCAness profiles (69) in future studies to shed light onto BRCA/HR-dependent or general responses to PARPi.

## Methods

### Cell culture experiments

Ovsaho cells were obtained from JCRB (Japanese Collection of Research Bioresources Cell Bank) cell bank (NIBIOHN, Cell No. JCRB 1046) and grown in MCDB105/199 medium with 10% FBS (Fisher, #10438026) and substituted with 1% Penicillin-Streptomycin. All cells were free of Mycoplasma and their identity was verified by whole exome sequencing at the Broad Institute. For dose-response curves, cells were treated for 72 hrs with DMSO or the indicated doses of the PARP1 inhibitor AG-14361 (Selleckchem, #S2178), FAS/FASN inhibitor TVB-2640 (Selleckchem, #S9714), Bortezomib (Selleckchem, #S1013), proTAME (Selleckchem, #S9605), GC7 (Sigma Aldrich #259545). In combination assays, inhibitor concentrations were combined in a 1:1 ratio for indicated total concentration. Cell viability was measured using the software in a live-imaging IncuCyte station by counting live cells. Live cells were labeled by adding the Incucyte NucLight Rapid Red (NRR) Reagent (Essen Bioscience, #4717, 1:4000) to the medium. Datapoints of three biological experiments and three technical replicates (total n=9) were merged and fitted using a non-linear log(inhibitor) versus normalized response with variable slope for dose-response curves using Prism (GraphPad). Combination Index (CI) was calculated based on Chou-Talaly method (53).

### Liquid chromatography-mass spectrometry (LC-MS) measurements

Tryptic digestion of proteins from samples were followed by a nanoflow LC/MS-MS analysis of peptides using the quadrupole Orbitrap mass spectrometer (Q Exactive HF-X, Thermo Fisher Scientific, Bremen, Germany) (Kelstrup et al. 2017) coupled to an EASYnLC 1200 ultra-high-pressure system (Thermo Fisher Scientific) through a nano-electrospray ion source. Briefly, about 300 ng peptides were loaded on a 50-cm HPLC-column with a 75-μm inner diameter (New Objective, USA; in-house packed with ReproSil-Pur C18-AQ 1.9-*μ*m silica beads; Dr. Maisch GmbH, Germany). Peptides were separated using a linear gradient from 5% to 60% solvent B (80% ACN; 0.1% formic acid) in solvent A (0.1% formic acid). Total gradient length was 100 min and the column temperature was maintained at 60 °C with an in-house made column oven. The. The MS was operated in a data-independent acquisition (DIA) mode. The DIA method consisted of one MS1 scan (m/z 350 to 1,650, resolution 60,000, maximum ion injection time 60 ms, AGC target 3E6) and 32 DIA segments with varying isolation widths ranging from 39.7 Th to 562.8 Th inversely proportional to the *m/z* distribution of tryptic peptides and with an overlap of 1 Th. MS2 scan resolution was set to 30,000, maximum injection time to 54 ms and the automated gain control (AGC) target to 3E6. A stepped collision energy of 25, 27.5 and 30 was used and the default MS2 charge state was 2. Data was acquired in profile mode.

### MS data analysis

DIA raw files were analyzed with Spectronaut Pulsar X software (Biognosys, version 12.0.20491.20) using the default settings for targeted data extraction. We used a project specific OVSAHO cell line proteome dataset as spectral library comprising 7,720 protein groups (103,554 precursors). The false discovery rates at the precursor and protein levels were < 1% with the ‘mutated’ decoy model. Prior to data analysis, data was filtered by ‘No Decoy’ and ‘Quantification Data Filtering’.

### Proteomics data analysis and computational modeling

Data analysis was done in the R software environment (Version 3.6.2). Variance between samples was tested using F-test. Statistical significance was assessed using moderated un-paired t-test comparing inhibitor-treated samples with control samples (DMSO was used as control). The p-values were corrected for an FDR of < 0.2 by the Benjamini–Hochberg method (30,31). Results were filtered to have both a significant FDR corrected p-value and a minimum log2(expression ratio) of 0.5. The NetboxR algorithm (28,29) was used to identify functional protein modules with FDR corrected p-value ≤ 0.05. For protein network analysis we used all proteins based on their variance level by selecting for proteins whose log2(expression ratio) was > 0.5 or <−0.5 and greater than technical noise, as defined by standard deviation of biological replicates. Networks were inferred based on the Reactome protein network (version 8), proteins are represented as nodes and interactions as edges. Statistical significance of discovered modules was assessed using the global network null model and local network null model with a p-value of ≤ 0.05. Linker p-value cut off of 0.05 was used to identify linker proteins (STable2). Derived network was illustrated using Cytoscape and Adobe Illustrator software. Based on the Reactome dataset, pathway enrichment analysis was performed using g:profiler p-value ≤ 0.05 (30). Activity scores of annotated protein modules were calculated as the sum of all expression change values in modules (17).

### L1000 dataset

The publicly available mRNA expression profiles of ovarian cancer cells from the L1000 Dataset -small molecule perturbagens- LINCS Phase 1 (50) (Todd Golub, Aravind Subramanian: L1000 Dataset -small molecule perturbagens- LINCS Phase 1, 2014, LINCS (collection), http://identifiers.org/lincs.data/LDG-1188, was retrieved on May 26, 2020 and data level 4 was used.

---

**SFigure 1.**
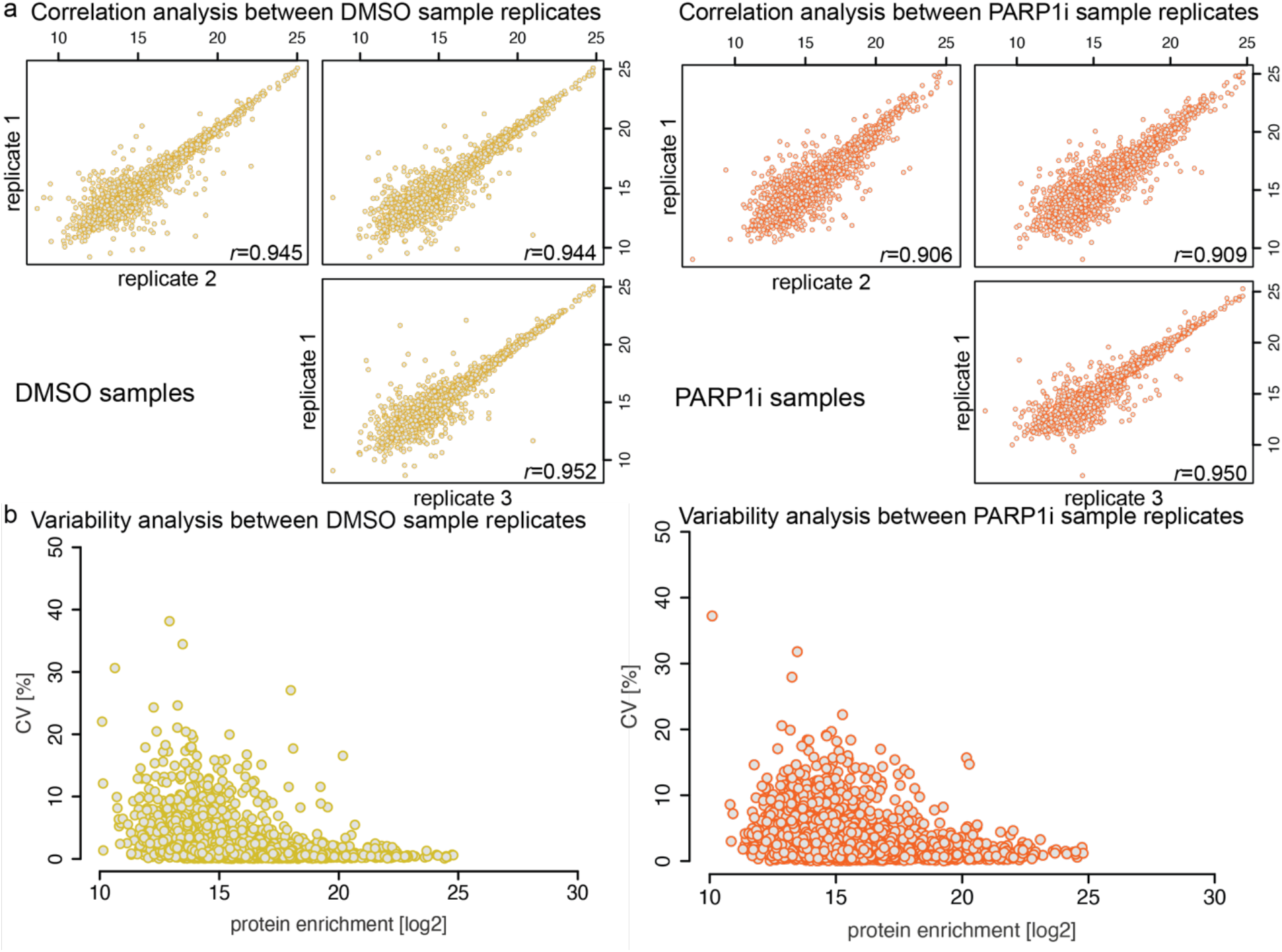
(a) Pearson correlation coefficients (*r*) were calculated for biological replicates (n=3) of samples. (b) Quantification of sample comparability determined by coefficient of variation (CV) of replicates.

**Supplementary Figure (SFigure) 2.**
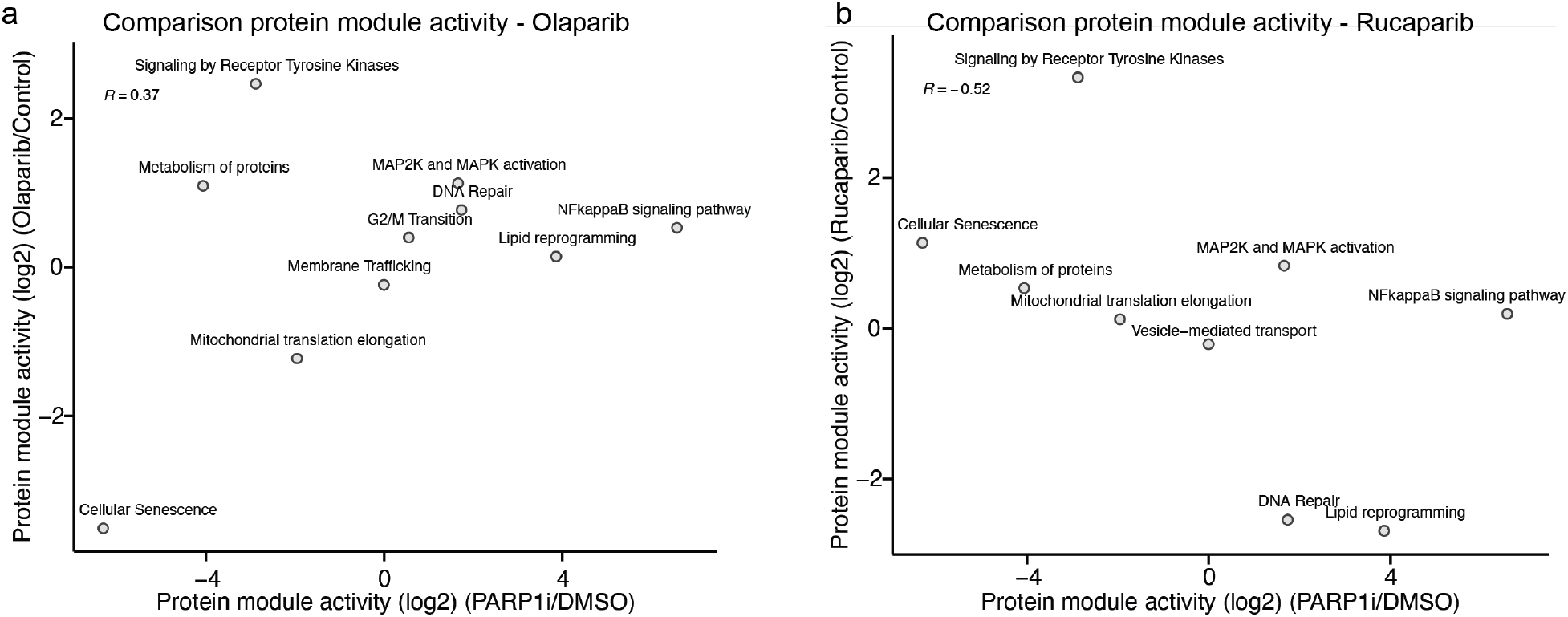
(a, b) Correlation analysis of the overall protein module activity based on overlapping annotated modules between L1000 olaparib-treated cell lines (a) and rucaparib-treated cell lines (b) and proteome profiling. Activity was calculated by integrating protein expression changes in modules (38) (Material and Methods).

## Author’s contributions

AF designed the study, collected the samples, analyzed and interpreted the proteomics data, performed combination dose-response experiments, analyzed live-cell imaging data and wrote the manuscript. FC provided advice for the design of the study, acquired and analyzed the mass spectrometry data and critically revised the paper. LC carried out the dose-response experiments, analyzed live-cell imaging data, prepared the samples for MS measurements and edited the manuscript. CYS participated in the design of the study, analyzed the WES data, performed combination dose-response experiments, analyzed live-cell imaging data and edited the paper. MM and CS designed and supervised the study and critically revised the paper. All authors have approved the final version.

## Acknowledgements

We thank Bo Yuan, Augustin Luna, Nicholas Gauthier, Frank Poelwijk, Michael Stiffler, Debora Marks, Ioannis Zervantonakis, Elizabeth Stover, Alan D’Andrea, Ernst Lengyel, Joan Brugge, and Ursula Matulonis for helpful discussions and advice.

## References

1. Prat J, FIGO Committee on Gynecologic Oncology. Staging classification for cancer of the ovary, fallopian tube, and peritoneum. Int J Gynecol Obstet. 2014 Jan;124(1):1–5.

2. Matulonis UA, Sood AK, Fallowfield L, Howitt BE, Sehouli J, Karlan BY. Ovarian cancer. Nat Rev Dis Primer. 2016 Dec;2(1):16061.

3. Ledermann J, Harter P, Gourley C, Friedlander M, Vergote I, Rustin G, et al. Olaparib Maintenance Therapy in Platinum-Sensitive Relapsed Ovarian Cancer. N Engl J Med. 2012 Apr 12;366(15):1382–92.

4. Pujade-Lauraine E, Ledermann JA, Selle F, Gebski V, Penson RT, Oza AM, et al. Olaparib tablets as maintenance therapy in patients with platinum-sensitive, relapsed ovarian cancer and a BRCA1/2 mutation (SOLO2/ENGOT-Ov21): a double-blind, randomised, placebo-controlled, phase 3 trial. Lancet Oncol. 2017;18(9):1274–84.

5. Kaufman B, Shapira-Frommer R, Schmutzler RK, Audeh MW, Friedlander M, Balmaña J, et al. Olaparib Monotherapy in Patients With Advanced Cancer and a Germline *BRCA1/2* Mutation. J Clin Oncol. 2015 Jan 20;33(3):244–50.

6. Swisher EM, Lin KK, Oza AM, Scott CL, Giordano H, Sun J, et al. Rucaparib in relapsed, platinum-sensitive high-grade ovarian carcinoma (ARIEL2 Part 1): an international, multicentre, open-label, phase 2 trial. Lancet Oncol. 2017 Jan;18(1):75–87.

7. Mirza MR, Monk BJ, Herrstedt J, Oza AM, Mahner S, Redondo A, et al. Niraparib Maintenance Therapy in Platinum-Sensitive, Recurrent Ovarian Cancer. N Engl J Med. 2016 Dec;375(22):2154–64.

8. Kristeleit R, Shapiro GI, Burris HA, Oza AM, LoRusso P, Patel MR, et al. A Phase I–II Study of the Oral PARP Inhibitor Rucaparib in Patients with Germline *BRCA1/2*-Mutated Ovarian Carcinoma or Other Solid Tumors. Clin Cancer Res. 2017 Aug 1;23(15):4095–106.

9. Kurnit KC, Fleming GF, Lengyel E. Updates and New Options in Advanced Epithelial Ovarian Cancer Treatment. Obstet Gynecol. 2021 Jan 1;137(1):108–21.

10. Kim D-S, Camacho CV, Kraus WL. Alternate therapeutic pathways for PARP inhibitors and potential mechanisms of resistance. Exp Mol Med. 2021 Jan;53(1):42–51.

11. Lord CJ, Ashworth A. PARP inhibitors: Synthetic lethality in the clinic. Science. 2017 Mar 17;355(6330):1152–8.

12. Helleday T. The underlying mechanism for the PARP and BRCA synthetic lethality: Clearing up the misunderstandings. Mol Oncol. 2011 Aug;5(4):387–93.

13. Murai J, Huang S -y. N, Das BB, Renaud A, Zhang Y, Doroshow JH, et al. Trapping of PARP1 and PARP2 by Clinical PARP Inhibitors. Cancer Res. 2012 Nov 1;72(21):5588–99.

14. Hopkins TA, Shi Y, Rodriguez LE, Solomon LR, Donawho CK, DiGiammarino EL, et al. Mechanistic Dissection of PARP1 Trapping and the Impact on In Vivo Tolerability and Efficacy of PARP Inhibitors. Mol Cancer Res. 2015 Nov 1;13(11):1465–77.

15. Liu C, Vyas A, Kassab MA, Singh AK, Yu X. The role of poly ADP-ribosylation in the first wave of DNA damage response. Nucleic Acids Res. 2017 Aug 21;45(14):8129–41.

16. Schiewer MJ, Knudsen KE. Transcriptional Roles of PARP1 in Cancer. Mol Cancer Res. 2014 Aug;12(8):1069–80.

17. Idogawa M, Yamada T, Honda K, Sato S, Imai K, Hirohashi S. Poly(ADP-Ribose) Polymerase-1 Is a Component of the Oncogenic T-Cell Factor-4/β-Catenin Complex. Gastroenterology. 2005 Jun;128(7):1919–36.

18. Santilli G, Cervellera MN, Johnson TK, Lewis RE, Iacobelli S, Sala A. PARP co-activates B-MYB through enhanced phosphorylation at cyclin/cdk2 sites. Oncogene. 2001 Dec;20(57):8167–74.

19. Elser M, Borsig L, Hassa PO, Erener S, Messner S, Valovka T, et al. Poly(ADP-Ribose) Polymerase 1 Promotes Tumor Cell Survival by Coactivating Hypoxia-Inducible Factor-1-Dependent Gene Expression. Mol Cancer Res. 2008 Feb 1;6(2):282–90.

20. Hassa PO, Covic M, Hasan S, Imhof R, Hottiger MO. The Enzymatic and DNA Binding Activity of PARP-1 Are Not Required for NF-ϰB Coactivator Function. J Biol Chem. 2001 Dec 7;276(49):45588–97.

21. González-Martín A, Pothuri B, Vergote I, DePont Christensen R, Graybill W, Mirza MR, et al. Niraparib in Patients with Newly Diagnosed Advanced Ovarian Cancer. N Engl J Med. 2019 Dec 19;381(25):2391–402.

22. Swisher EM, Kwan TT, Oza AM, Tinker AV, Ray-Coquard I, Oaknin A, et al. Molecular and clinical determinants of response and resistance to rucaparib for recurrent ovarian cancer treatment in ARIEL2 (Parts 1 and 2). Nat Commun. 2021 Dec;12(1):2487.

23. Konstantinopoulos PA, Ceccaldi R, Shapiro GI, D’Andrea AD. Homologous Recombination Deficiency: Exploiting the Fundamental Vulnerability of Ovarian Cancer. Cancer Discov. 2015 Nov;5(11):1137–54.

24. Zhao W, Li J, Chen M-JM, Luo Y, Ju Z, Nesser NK, et al. Large-Scale Characterization of Drug Responses of Clinically Relevant Proteins in Cancer Cell Lines. Cancer Cell. 2020 Nov;S1535610820305390.

25. Molinelli EJ, Korkut A, Wang W, Miller ML, Gauthier NP, Jing X, et al. Perturbation biology: inferring signaling networks in cellular systems. PLoS Comput Biol. 2013;9(12):e1003290.

26. Calabrese CR, Almassy R, Barton S, Batey MA, Calvert AH, Canan-Koch S, et al. Anticancer Chemosensitization and Radiosensitization by the Novel Poly(ADP-ribose) Polymerase-1 Inhibitor AG14361. JNCI J Natl Cancer Inst. 2004 Jan 7;96(1):56–67.

27. Domcke S, Sinha R, Levine DA, Sander C, Schultz N. Evaluating cell lines as tumour models by comparison of genomic profiles. Nat Commun. 2013 Oct;4(1):2126.

28. Coscia F, Watters KM, Curtis M, Eckert MA, Chiang CY, Tyanova S, et al. Integrative proteomic profiling of ovarian cancer cell lines reveals precursor cell associated proteins and functional status. Nat Commun. 2016 Nov;7(1):12645.

29. Murina O, von Aesch C, Karakus U, Ferretti LP, Bolck HA, Hänggi K, et al. FANCD2 and CtIP Cooperate to Repair DNA Interstrand Crosslinks. Cell Rep. 2014 May;7(4):1030–8.

30. Huttenhain R, Soste M, Selevsek N, Rost H, Sethi A, Carapito C, et al. Reproducible Quantification of Cancer-Associated Proteins in Body Fluids Using Targeted Proteomics. Sci Transl Med. 2012 Jul 11;4(142):142ra94–142ra94.

31. Schmidt A, Beck M, Malmström J, Lam H, Claassen M, Campbell D, et al. Absolute quantification of microbial proteomes at different states by directed mass spectrometry. Mol Syst Biol. 2011 Jan;7(1):510.

32. Vellingiri, Iyer, Devi Subramaniam, Jayaramayya, Siama, Giridharan, et al. Understanding the Role of the Transcription Factor Sp1 in Ovarian Cancer: from Theory to Practice. Int J Mol Sci. 2020 Feb 9;21(3):1153.

33. Knudsen ES, Pruitt SC, Hershberger PA, Witkiewicz AK, Goodrich DW. Cell Cycle and Beyond: Exploiting New RB1 Controlled Mechanisms for Cancer Therapy. Trends Cancer. 2019 May;5(5):308–24.

34. Lamb J, Ramaswamy S, Ford HL, Contreras B, Martinez RV, Kittrell FS, et al. A mechanism of cyclin D1 action encoded in the patterns of gene expression in human cancer. Cell. 2003 Aug 8;114(3):323–34.

35. Williams SR, Zies D, Mullegama SV, Grotewiel MS, Elsea SH. Smith-Magenis syndrome results in disruption of CLOCK gene transcription and reveals an integral role for RAI1 in the maintenance of circadian rhythmicity. Am J Hum Genet. 2012 Jun 8;90(6):941–9.

36. van de Wetering M, Sancho E, Verweij C, de Lau W, Oving I, Hurlstone A, et al. The beta-catenin/TCF-4 complex imposes a crypt progenitor phenotype on colorectal cancer cells. Cell. 2002 Oct 18;111(2):241–50.

37. Willis TG, Jadayel DM, Du MQ, Peng H, Perry AR, Abdul-Rauf M, et al. Bcl10 is involved in t(1;14)(p22;q32) of MALT B cell lymphoma and mutated in multiple tumor types. Cell. 1999 Jan 8;96(1):35–45.

38. Ortiz-Zapater E, Pineda D, Martínez-Bosch N, Fernández-Miranda G, Iglesias M, Alameda F, et al. Key contribution of CPEB4-mediated translational control to cancer progression. Nat Med. 2012 Jan;18(1):83–90.

39. Cerami E, Demir E, Schultz N, Taylor BS, Sander C. Automated Network Analysis Identifies Core Pathways in Glioblastoma. Hannenhalli S, editor. PLoS ONE. 2010 Feb 12;5(2):e8918.

40. Liu EM, Luna A, Dong G, Sander C. netboxr: Automated discovery of biological process modules by network analysis in R. PloS One. 2020;15(11):e0234669.

41. Reimand J, Kull M, Peterson H, Hansen J, Vilo J. g:Profiler—a web-based toolset for functional profiling of gene lists from large-scale experiments. Nucleic Acids Res. 2007 Jul;35(suppl_2):W193–200.

42. Fleury H, Malaquin N, Tu V, Gilbert S, Martinez A, Olivier M-A, et al. Exploiting interconnected synthetic lethal interactions between PARP inhibition and cancer cell reversible senescence. Nat Commun. 2019 Dec;10(1):2556.

43. Fang Y, McGrail DJ, Sun C, Labrie M, Chen X, Zhang D, et al. Sequential Therapy with PARP and WEE1 Inhibitors Minimizes Toxicity while Maintaining Efficacy. Cancer Cell. 2019 Jun;35(6):851–867.e7.

44. Momeny M, Yousefi H, Eyvani H, Moghaddaskho F, Salehi A, Esmaeili F, et al. Blockade of nuclear factor-ϰB (NF-ϰB) pathway inhibits growth and induces apoptosis in chemoresistant ovarian carcinoma cells. Int J Biochem Cell Biol. 2018 Jun;99:1–9.

45. Darvishi B, Farahmand L, Eslami-S Z, Majidzadeh-A K. NF-ϰB as the main node of resistance to receptor tyrosine kinase inhibitors in triple-negative breast cancer. Tumor Biol. 2017 Jun;39(6):101042831770691.

46. Seubwai W, Vaeteewoottacharn K, Kraiklang R, Umezawa K, Okada S, Wongkham S. Inhibition of NF-ϰB Activity Enhances Sensitivity to Anticancer Drugs in Cholangiocarcinoma Cells. Oncol Res Featur Preclin Clin Cancer Ther. 2016 Jan 21;23(1):21–8.

47. Nakagawa Y, Sedukhina AS, Okamoto N, Nagasawa S, Suzuki N, Ohta T, et al. NF-ϰB signaling mediates acquired resistance after PARP inhibition. Oncotarget. 2015 Feb 28;6(6):3825–39.

48. Ji Z, Shen Y, Feng X, Kong Y, Shao Y, Meng J, et al. Deregulation of Lipid Metabolism: The Critical Factors in Ovarian Cancer. Front Oncol. 2020 Oct 19;10:593017.

49. Niepel M, Hafner M, Duan Q, Wang Z, Paull EO, Chung M, et al. Common and cell-type specific responses to anti-cancer drugs revealed by high throughput transcript profiling. Nat Commun. 2017 Dec;8(1):1186.

50. Knezevic CE, Wright G, Remsing Rix LL, Kim W, Kuenzi BM, Luo Y, et al. Proteome-wide Profiling of Clinical PARP Inhibitors Reveals Compound-Specific Secondary Targets. Cell Chem Biol. 2016 Dec;23(12):1490–503.

51. Caraglia M, Park MH, Wolff EC, Marra M, Abbruzzese A. eIF5A isoforms and cancer: two brothers for two functions? Amino Acids. 2013 Jan;44(1):103–9.

52. Liess AKL, Kucerova A, Schweimer K, Schlesinger D, Dybkov O, Urlaub H, et al. Dimerization regulates the human APC/C-associated ubiquitin-conjugating enzyme UBE2S. Sci Signal. 2020 Oct 20;13(654):eaba8208.

53. Chou T-C. Drug Combination Studies and Their Synergy Quantification Using the Chou-Talalay Method. Cancer Res. 2010 Jan 15;70(2):440–6.

54. Raab M, Sanhaji M, Zhou S, Rödel F, El-Balat A, Becker S, et al. Blocking Mitotic Exit of Ovarian Cancer Cells by Pharmaceutical Inhibition of the Anaphase-Promoting Complex Reduces Chromosomal Instability. Neoplasia. 2019 Apr;21(4):363–75.

55. Gupta SC, Sundaram C, Reuter S, Aggarwal BB. Inhibiting NF-ϰB activation by small molecules as a therapeutic strategy. Biochim Biophys Acta BBA - Gene Regul Mech. 2010 Oct;1799(10–12):775–87.

56. Patel M, Infante J, Von Hoff D, Jones S, Burris H, Brenner A, et al. Abstract CT203: Report of a first-in-human study of the first-in-class fatty acid synthase (FASN) inhibitor TVB-2640. In: Clinical Trials [Internet]. American Association for Cancer Research; 2015 [cited 2021 May 24]. p. CT203–CT203. Available from: http://cancerres.aacrjournals.org/lookup/doi/10.1158/1538-7445.AM2015-CT203

57. Kraus WL. Transcriptional control by PARP-1: chromatin modulation, enhancer-binding, coregulation, and insulation. Curr Opin Cell Biol. 2008 Jun;20(3):294–302.

58. Ledermann J, Harter P, Gourley C, Friedlander M, Vergote I, Rustin G, et al. Olaparib maintenance therapy in patients with platinum-sensitive relapsed serous ovarian cancer: a preplanned retrospective analysis of outcomes by BRCA status in a randomised phase 2 trial. Lancet Oncol. 2014 Jul;15(8):852–61.

59. Gogola E, Rottenberg S, Jonkers J. Resistance to PARP Inhibitors: Lessons from Preclinical Models of BRCA-Associated Cancer. Annu Rev Cancer Biol. 2019 Mar 4;3(1):235–54.

60. Pyragius CE, Fuller M, Ricciardelli C, Oehler MK. Aberrant lipid metabolism: an emerging diagnostic and therapeutic target in ovarian cancer. Int J Mol Sci. 2013 Apr 10;14(4):7742–56.

61. Germain N, Dhayer M, Boileau M, Fovez Q, Kluza J, Marchetti P. Lipid Metabolism and Resistance to Anticancer Treatment. Biology. 2020 Dec 16;9(12):474.

62. Broadfield LA, Pane AA, Talebi A, Swinnen JV, Fendt S-M. Lipid metabolism in cancer: New perspectives and emerging mechanisms. Dev Cell. 2021 May;56(10):1363–93.

63. Curtarello M, Tognon M, Venturoli C, Silic-Benussi M, Grassi A, Verza M, et al. Rewiring of Lipid Metabolism and Storage in Ovarian Cancer Cells after Anti-VEGF Therapy. Cells. 2019 Dec 9;8(12):1601.

64. Chen RR, Yung MMH, Xuan Y, Zhan S, Leung LL, Liang RR, et al. Targeting of lipid metabolism with a metabolic inhibitor cocktail eradicates peritoneal metastases in ovarian cancer cells. Commun Biol. 2019 Dec;2(1):281.

65. Fhu CW, Ali A. Fatty Acid Synthase: An Emerging Target in Cancer. Molecules. 2020 Aug 28;25(17):3935.

66. ,, ,, Berkel Ç, Küçük B, Usta M, Yılmaz E, Çaçan E. The Effect of Olaparib and Bortezomib Combination Treatment on Ovarian Cancer Cell Lines. Eur J Biol [Internet]. 2020 Jan 4 [cited 2021 May 17]; Available from: https://iupress.istanbul.edu.tr/tr/journal/ejb/article/the-effect-of-olaparib-and-bortezomib-combination-treatment-on-ovarian-cancer-cell-lines

67. Palmer AC, Sorger PK. Combination Cancer Therapy Can Confer Benefit via Patient-to-Patient Variability without Drug Additivity or Synergy. Cell. 2017 Dec;171(7):1678–1691.e13.

68. Palmer AC, Chidley C, Sorger PK. A curative combination cancer therapy achieves high fractional cell killing through low cross-resistance and drug additivity. eLife. 2019 Nov 19;8:e50036.

69. Turner N, Tutt A, Ashworth A. Hallmarks of “BRCAness” in sporadic cancers. Nat Rev Cancer. 2004 Oct;4(10):814–9.

